# Amyloidogenic core of a human λ-III immunoglobulin light chain fibril and their germline variants probed by MAS solid state NMR

**DOI:** 10.1101/2020.10.02.323303

**Authors:** Tejaswini Pradhan, Karthikeyan Annamalai, Riddhiman Sarkar, Stephanie Huhn, Ute Hegenbart, Stefan Schönland, Marcus Fändrich, Bernd Reif

**Affiliations:** Helmholtz-Zentrum München (HMGU), Deutsches Forschungszentrum für Gesundheit und Umwelt, Institute of Structural Biology (STB), Ingolstädter Landstr. 1, 85764 Neuherberg, Germany; Munich Center for Integrated Protein Science (CIPS-M) at the Department of Chemistry, Technische Universität München (TUM), Lichtenbergstr. 4, 85747 Garching, Germany; Institute of Protein Biochemistry, Ulm University, Helmholtzstr. 8/1, 89081 Ulm, Germany; Medical Department V, Amyloidosis Center, Heidelberg University Hospital, 69120 Heidelberg, Germany

**Keywords:** AL amyloidosis, antibody light chain, protein aggregation, fibril seeding, Magic Angle Spinning (MAS) solid-state NMR spectroscopy

## Abstract

Systemic antibody light chains (AL) amyloidosis is characterized by deposition of amyloid fibrils derived from a particular antibody light chain. Cardiac involvement is a major risk factor for mortality. Using MAS solid-state NMR, we study the fibril structure of a recombinant light chain fragment corresponding to the fibril protein from patient FOR005, together with fibrils formed by protein sequence variants that reflect the closest germline (GL) sequence. Both analyzed fibril structures were seeded with *ex-vivo* amyloid fibrils purified from the explanted heart of this patient. We find that residues 11-42 and 69-102 adopt β-sheet conformation in patient protein fibrils. We identify glycine-49 that is mutated with respect to the germline sequence into arginine-49 as a key residue that forms a salt bridge to aspartate-25 in the patient protein fibril structure. Fibrils from the GL protein and from the patient protein harboring the single point mutation R49G can be both heterologously seeded using patient *ex-vivo* fibrils. Seeded R49G fibrils show an increased heterogeneity for the C-terminal residues 80-102 which is reflected by the disappearance of all resonances of these residues. By contrast, residues 11-42 and 69-77, which are visible in the MAS solid-state NMR spectra show ^13^Cα chemical shifts that are highly similar to patient fibrils. The mutation R49G thus induces a conformational heterogeneity at the C-terminus in the fibril state, while the overall fibril topology is retained.

## Introduction

Antibody light chain (AL) amyloidosis is a rare disease affecting about 9-14 new cases per one million inhabitants per year (1). The disease is caused by formation of amyloid fibrils from immunoglobulin light chains (LCs) (2–4). An underlying plasma cell dyscrasia causes overproduction and secretion of a monoclonal LC. Some multiple myeloma patients develop AL amyloidosis as a secondary disease, in which LCs can assemble via oligomeric intermediates into fibrils which deposit in the inner organs, such as heart and kidney. Heart involvement is a major risk factor of mortality and the survival rate is on the order of 7 month in patients with advanced cardiac amyloidosis (5). Due to recombination and somatic hypermutation of LC gene segments, the potential number of sequence variants of antibody LCs is enormous (6). This variability makes the identification of aggregation hotspots within a particular sequence a challenging undertaking (7–9). It is known that the λ-III LCs are overrepresented in AL amyloidosis (10), together with the LC subtypes λ-VI, λ-I, λ-II, κ-I (6,8). In AL patients, λ and κ isotypes occur at a ratio of λ:κ = 3:1, while for healthy or multiple myeloma patients the ratio is rather λ:κ = 1:2 (11). It is of fundamental importance to identify the sequence elements or residues that are causative for fibril formation and stability (7,8). However, a correlation between sequence and amyloidogenicity remains elusive.

All antibody LCs consist of a variable light (V_L_) and a constant light (C_L_) domain which both adopt an immunoglobulin fold. This structural conservation in the LC native state raises the question whether AL fibrils adopt also a common amyloid structure in the aggregated state. So far, only two structures have been determined using cryo-EM (12,13), that differ significantly from the native LC fold as well as from one another, indicating that multiple fibril topologies can be involved in this disease. Using magic angle spinning (MAS) solid-state NMR, chemical shift assignments from three additional LC derived fibrils have been reported so far, including fibrils of the murine κ-IV MAK33 (14), the patient derived κ-I LC AL09 (15,16), and the λ-VI model germline LC 6aJL2_R25G (17). All fibril structures differ profoundly with respect to their amyloidogenic cores.

We focus here on the structural characterization of recombinant FOR005 protein fibrils using primarily MAS solid-state NMR. Patient FOR005 showed a dominant heart involvement (18). while the presently analyzed protein corresponds to the main fibril protein in this patient. This protein is a fragment of its LC precursor and almost identical to the V_L_ domain. Refolded from the heart it crystallizes as a dimer with a canonical dimer interface (18,19), while it is mainly monomeric in solution. The fibril protein contains the disulphide bond that is already present in the native LC with no other post-translational modifications within the VL domain. We used fibrils extracted from patient tissue to seed the formation of fibrils *in vitro,* aiming to imprint the patient fibril structure onto the *in-vitro* prepared protein. We assigned the core of the fibrils, and identified a number of electrostatic interactions in the fibril core that may be important for fibril stability. In addition, we investigated fibrils formed by the germline (GL) sequence, as well as of patient protein harboring the single point mutation R49G. We find that both FOR005-R49G and GL fibrils can be seeded using *ex vivo* material, and adopt a similar conformation as patient fibrils. The spectroscopic results are discussed to address the role of mutations and cross-seeding on the conformation and stability of an amyloid fibril.

## Results and Discussion

The primary structure of the fibril protein precursor of FOR005, a λ-III LC, was obtained previously by cDNA sequencing (18). For reference, we determined the respective GL sequence (FOR005_GL), using the web tools abYsis (http://www.abysis.org/) and IMGT (http://www.imgt.org/). Consistent with previous analyses of GL sequences (20,21), we assumed that the germline sequence of FOR005 has a lower aggregation propensity compared to the patient sequence. FOR005 and FOR005_GL differ in five amino acids in the variable GL segment, namely at residues S31Y, F48Y, R49G, S51N and A94G (mutations indicate transitions from patient to GL protein). All mutations are located within, or near to the hypervariable complementarity determining regions (CDRs). In addition to the GL protein, we analyzed the fibrils formed by the recombinant patient protein FOR005 as well as by the patient protein carrying the single point mutation R49G. This mutation is the least conservative mutation, and we find it to be particularly important for fibril formation and stability (see below). All proteins were recombinantly expressed and purified, as described in the Materials section. We employed MAS solid state NMR spectroscopy, thioflavin T (ThT) fluorescence, circular dichroism (CD) spectroscopy and transmission electron microscopy (TEM) to characterize the aggregation properties of the soluble LC protein, its thermodynamic properties, as well as its structure in the fibril state.

### The fibril core of FOR005

For solid-state NMR, polymorphism in fibril sample preparations is a severe obstacle, as it results in sample heterogeneity and in the loss of spectral resolution. At the same time, reproducibility of fibril growth impedes a more detailed structural analysis and prevents the derivation of general principles. To overcome this problem, seeds are employed to prepare homogeneous fibrils (22). Seeding with *ex-vivo* fibrils results in a reduction of the lag phase of the fibril kinetics and allows us to obtain highly reproducible NMR spectra (22). This finding is in agreement with previous observations made for different amyloid preparations investigated by MAS solid-state NMR (14,23–25). TEM experiments (Figure 1A) show relatively homogeneous fibrils with no indication of polymorphism.

**Figure 1.**
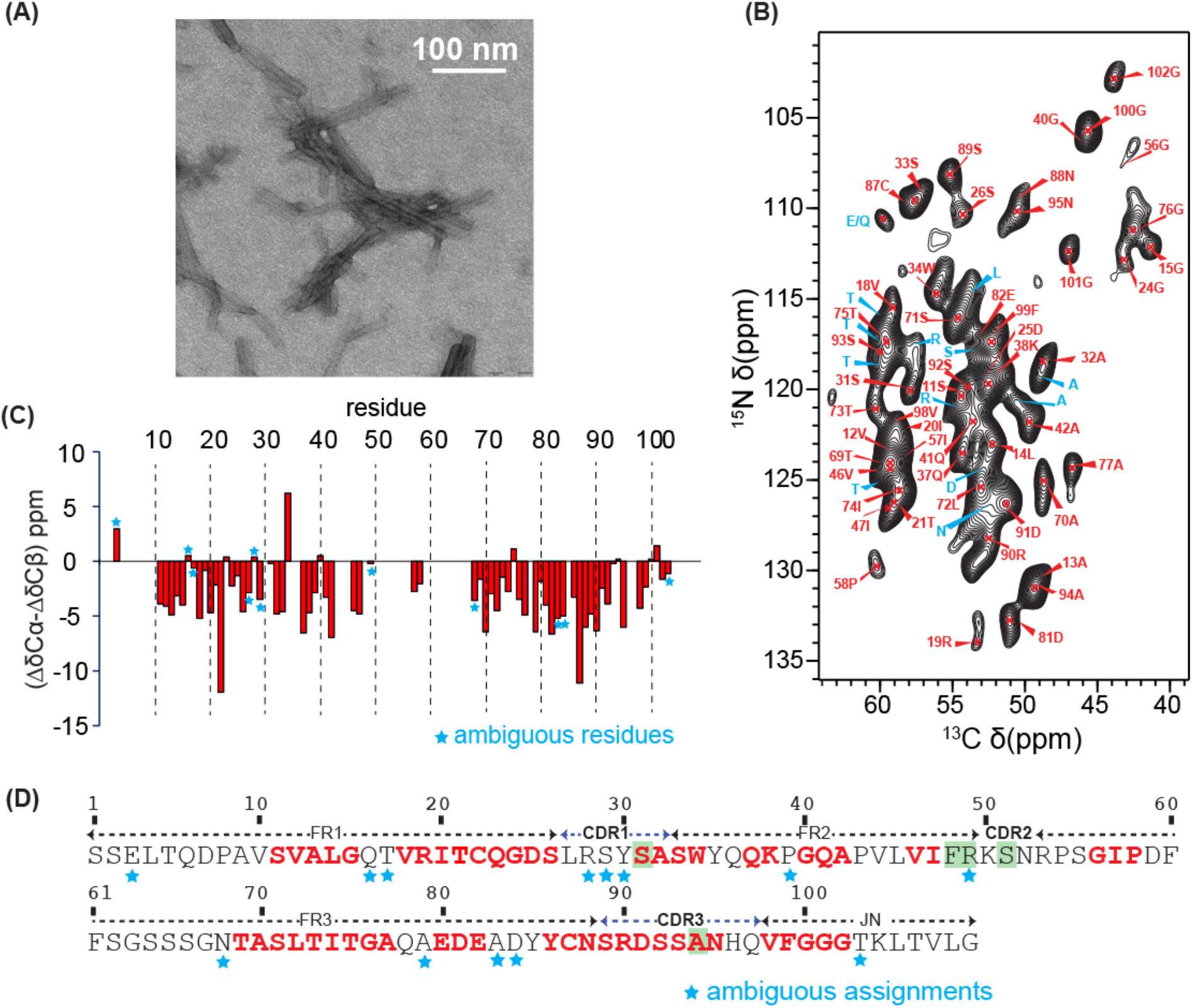
MAS solid-state NMR investigations of fibrils formed from recombinant FOR005 fibril protein and seeded with ex-vivo fibrils. (A) TEM image of seeded FOR005 fibrils. (B) 2D ^13^C,^15^N correlation spectrum with a focus on the Cα spectral region. (C) Secondary chemical shifts as a function of the residue. For glycines, only the ^13^Cα chemical shift values were considered. Asterisks indicate residues for which only amino acid type assignments are available. (D) Primary structure of the expressed fibril protein of patient FOR005. Assigned residues are color coded in red. Residues marked in green represent the five mutations with respect to the closest germline sequence. Asterisks again indicate residues for which only amino acid type assignments are available.

For solid-state NMR experiments, we prepared in total seven fibril samples. Two samples were identical replicates containing recombinant FOR005 fibrils, using *ex-vivo* fibrils as seeds to confirm that spectra are reproducible (22). Two fibril samples were prepared from FOR005 protein grown in the presence of *in-vitro* seeds (22) as well as without seeding. Furthermore, we produced fibrils of the germline protein (GL) employing *ex-vivo* seeds, as well as a sample of fibrils formed by the patient protein carrying the single point mutation R49G (FOR005_R49G) using *ex-vivo* seeds. Expect from the employed protein and seeds, the preparation conditions for each sample were identical. All samples were uniformly ^13^C,^15^N isotopically enriched and were packed after fibril growth into a 3.2 mm MAS rotor (see Materials).

The assigned 2D ^13^C,^15^N correlation spectrum of fibrils grown from FOR005 fibril protein seeded with *ex-vivo* fibrils is shown in Figure 1B. The sequential assignment of this sample yields one set of resonances. None of the recorded spectra shows any indication of structural heterogeneity or polymorphism of the fibril sample. The NMR chemical shift differences (ΔδCα-ΔδCβ) which are indicative for secondary structure (26) suggest that virtually all assigned residues adopt β-sheet conformation (Figure 1C). The first 12 N-terminal residues, the 7 last C-terminal residues, as well as residues 48-68 are not observable and cannot be assigned (Figure 1D). We therefore conclude that these residues may not be part of the fibril core and conformationally heterogeneous or disordered. Out of the five residues which are mutated in FOR005 with respect to the germline sequence, only residues S31 and A94 which are located near CDR1 and in CDR3, respectively, could be assigned to the fibril core.

### Heterologous seeding

Previous studies have shown that seeds with a substantially different primary sequence are able to accelerate fibril formation of non-homologous proteins (27). We therefore wanted to test whether seeding is effective as well for the different germline variant proteins. Using *ex-vivo* seeds, we find that fibril formation is significantly accelerated with a reduced lag phase for the V_L_ domains of the patient LC protein FOR005, of the single point mutant R49G and of the germline LC protein (Figure 2A).

**Figure 2.**
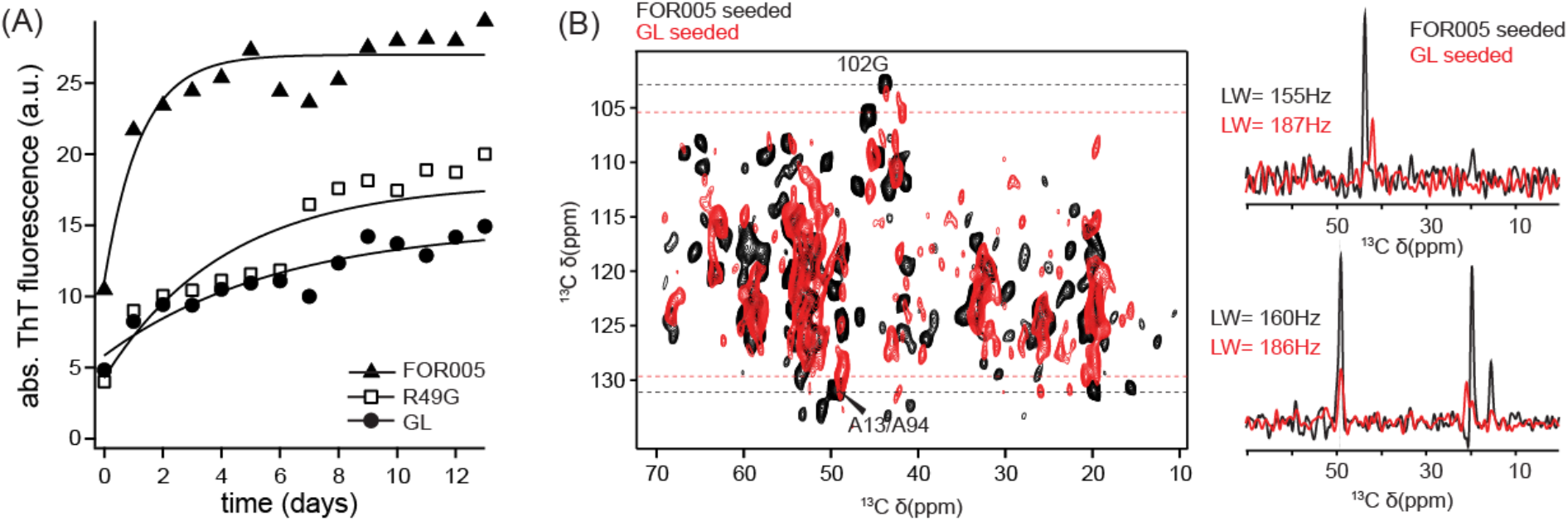
Biophysical characterization of FOR005 V_L_ variants. (A) Seeded ThT aggregation kinetics of patient protein FOR005, protein containing the single point mutation R49G and for protein coding for the germline sequence. In all cases, 5 % seeds were added to the monomeric protein and incubated at 37°C. B) Comparison of germline and patient fibrils both seeded with ex-vivo seeds. Superposition of the 2D NCACX correlation spectra for patient (black) and germline (red) fibrils. In both cases, ex-vivo seeds have been employed. For G102 and A13/A94, 1D traces were extracted along the ^13^C dimension (right). Fibrils formed by the germline protein show significantly reduced sensitivity and have increased line width (FWHM = 187 Hz for germline, FWHM = 155 Hz for patient fibrils).

The catalyzed conversion into fibrils suggests that the monomeric germline and the mutant R49G protein are recruited into fibrils by seeding. To validate this hypothesis further, we carried out MAS solid-state NMR experiments using fibrils formed by germline protein. Seeded germline fibrils yield spectra with a significantly decreased sensitivity (Figure 2B). Germline fibrils seem overall less homogeneous. Aggregation is quantitative in both cases and no protein is left in the supernatant after sedimentation of the fibril preparation into the MAS rotor. Nevertheless, clear spectral patterns can be recognized indicating that a fraction of the protein adopts a preferred conformation. The linewidth of ^13^C and ^15^N resonances of germline fibrils are comparable to the linewidth observed in the spectra of fibrils from FOR005 fibril protein, indicating a similar degree of order for the two fibril preparations.

Next, we wanted to identify the residues that stabilize the fibril fold. We therefore performed 2D ^13^C,^15^N Transferred Echo Double resonance (TEDOR) experiments using fibrils prepared from FOR005 patient protein (Figure 3A). Long range contacts between isolated spins were previously found to be important restraints to characterize the fold of an amyloid (28,29). Lysine and arginine side chains yield distinct peaks in 2D ^15^N,^13^C correlation spectra with ^15^N chemical shifts of 80 ppm (for Arg Nε) and 30 ppm (for Lys Nζ), respectively. The primary structure of FOR005 contains three lysines, that are all visible in the spectra (Figure 3A). K38 is sequentially assigned and observable in the 2D NCACX experiment (red contours in Figure 3A). If the mixing time of the TEDOR experiment is increased to 15 ms (cyan contour lines), additional cross peaks become visible at ^13^C chemical shifts of around 180 ppm corresponding to carboxylic acid groups of aspartate or glutamate. We assigned the long-range cross peak at a ^15^N chemical shift of around 30 ppm to an electrostatic interaction between the amino group of K50 and the carboxylic acid group of D81. The Nζ chemical shift of K38 is not in agreement with the observed long-range TEDOR cross peak, and is therefore excluded. Three out of six arginine residues are observed using a short TEDOR contact time of 1.9 ms. The long-range cross peak involving an arginine guanidine group is assigned to a salt bridge between R49 and the carboxylic acid group of D25. The assignment of R49 is confirmed by mutagenesis (see below).

**Figure 3.**
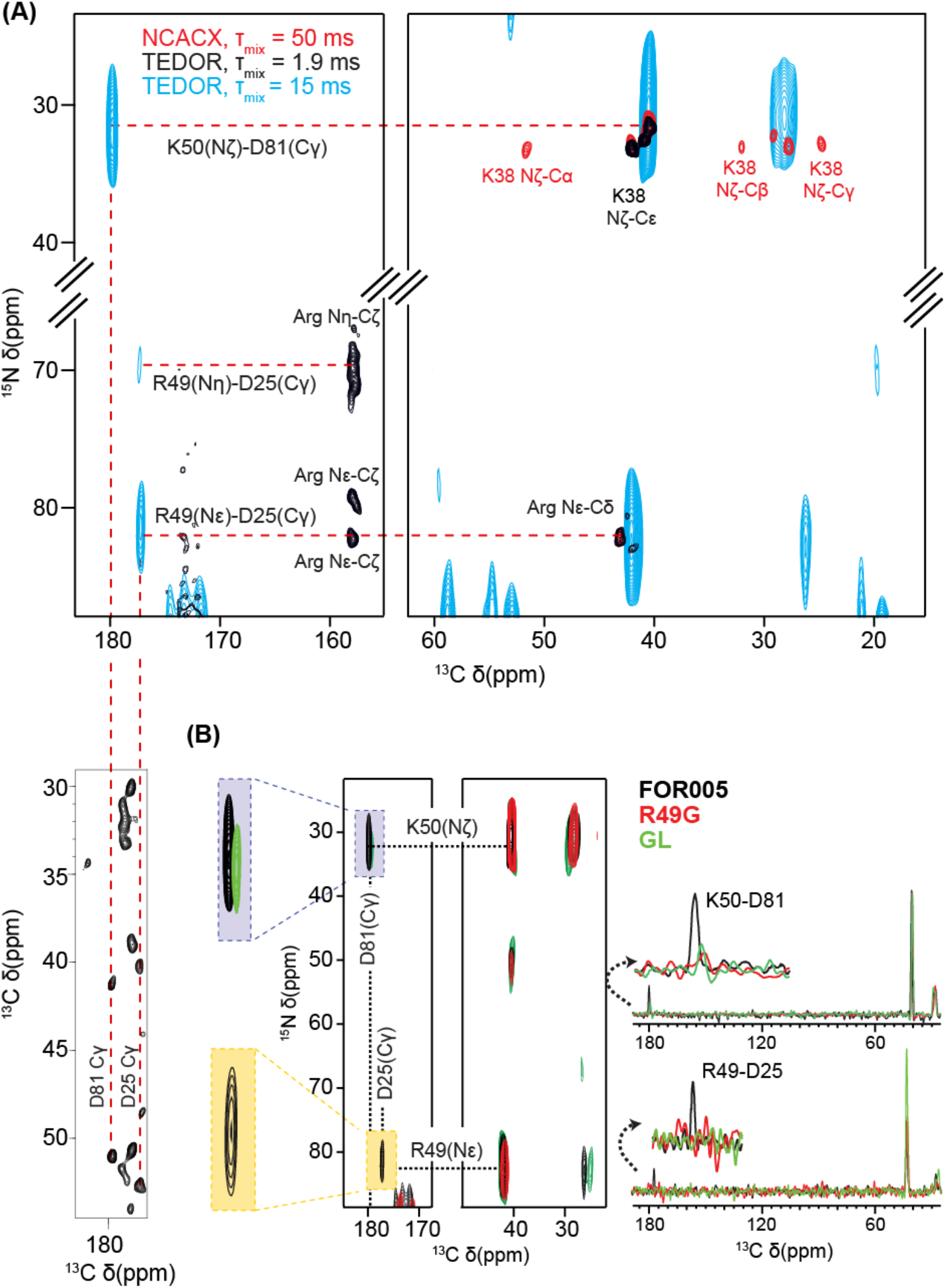
Electrostatic interactions between charged side chains in FOR005 patient, germline and R49G fibrils probed by TEDOR experiments. (A) FOR005 patient fibrils. Superposition of 2D TEDOR spectra obtained with short (black, ⊺_mix_= 1.9 ms) and long mixing times (cyan, ⊺_mix_= 15 ms), focusing on the side chain resonances of arginine and lysine. The spectra are superimposed onto a 2D ^13^C,^15^N NCACX spectrum (red, ⊺_mix_= 50 ms). We observe long-range contacts between lysine/arginine and aspartic acid side chains. The assignment of the carboxylic acid groups is indicated with a red dashed line in the 2D ^13^C,^13^C PDSD spectrum. (B) Comparison of 2D TEDOR fibril spectra for FOR005 sequence variants. Patient fibril spectra are represented in black, germline fibrils in green and R49G fibrils in red. On the right, 1D rows extracted from the 2D TEDOR experiments are shown, illustrating the peak intensities for the different protein variants. In all cases, experiments were recorded under identical conditions with an equal number of scans and increments. Apparently, only fibrils formed by the patient protein FOR005 contain the salt bridges, while no cross peaks are observed in R49G. For germline fibrils, a very weak peak seems to indicate a strongly reduced interaction involving K50-D81.

The observed salt bridges should have a large stabilizing effect on the fibril structure. The sensitivity of GL fibrils was generally too low to pursue chemical shift assignments and a more detailed structural analysis (Figure 2B). From the five mutations from patient to germline (S31Y, F48Y, R49G, S51N and A94G), R49G is the least conservative mutation. We therefore decided to introduce the single point mutation R49G to study the effects that are induced by this particular side chain. This is supported by the finding that R49G has a large effect on the thermodynamic stability of the immunoglobulin fold (data not shown). In fact, TEDOR experiments for germline or R49G fibrils do not yield any long-range cross peaks (Figure 3B), suggesting that the salt bridge involving the guanidine group is due to R49. The R49-D25 cross peak is lost in both preparations, while the interaction K50-D81 is weak in germline and absent in R49G fibrils.

To more closely analyze the structural changes between FOR005 and R49G fibrils, we assigned the chemical shifts of the R49G fibril sample (Figure 4). The assignment of the core residues of R49G fibrils was achieved using a 3D NCACX experiment. The obtained chemical shifts were compared with the assignments of FOR005 fibrils. To confirm the assignments, a 2D NCOCX experiment was analyzed. We observe the largest chemical shift differences for residues 11-42 and 69-77 (Figure 4D). This is consistent with the picture that both salt bridges are lost in R49G fibrils. Figure 4E shows a correlation for the Cα chemical shift between patient fibrils and fibrils formed by the single point mutant protein R49G. The R-value is on the order of 0.76, indicating that the fibril structures of FOR005 and R49G are in fact rather similar. In addition, we find that the C-terminal residues are not any more visible in the spectra recorded for R49G fibrils, indicating that this part of the protein becomes conformationally disordered or dynamic in R49G fibrils. Surprisingly, we also find that the C-terminus of FOR005 fibrils is missing in the spectra of the non-seeded FOR005 fibril preparation (Figure 4F). This suggests that the template is equally important as the amyloid substrate to enable the formation of a stable fibril structure.

**Figure 4.**
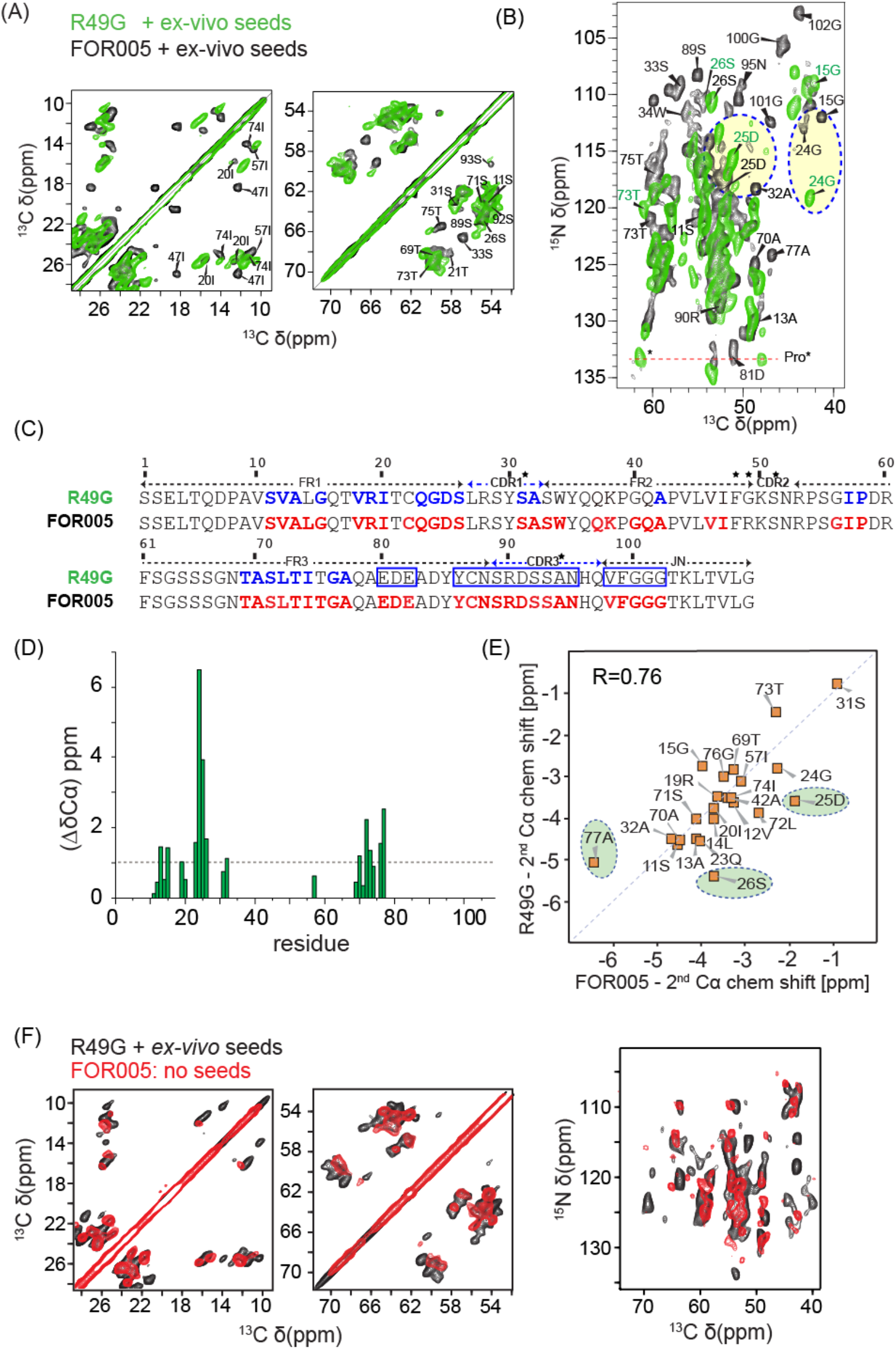
Comparison of fibril spectra obtained from FOR005 patient protein and FOR005 R49G. (A, B) Superposition of 2D ^13^C,^13^C and ^13^C,^15^N correlation spectra obtained for FOR005 and R49G fibrils. Residues from the C-terminal part of the protein are not observed in R49G. The yellow circles indicate residues that undergo large chemical shift changes. (C) Assigned residues in FOR005 and R49G are indicated in red and blue, respectively. Blue boxes highlight the residues that are not observed in fibrils of the point mutant R49G. Residues marked with an asterisk indicate mutations from germline to patient. (D) Cα chemical shift differences for FOR005 and R49G fibrils as a function of residue. (E) Correlation plot of the secondary Cα chemical shift (experimental shifts-random coil shifts) for patient FOR005 and the point mutation R49G. Correlations highlighted with a green circle represent residues that are close to the salt bridge R49-D25. (F) Comparison of non-seeded FOR005 and seeded R49G fibrils. The spectra are almost identical, suggesting that the fibril structures are similar. (Left) 2D PDSD ^13^C,^13^C correlation spectra, focusing on the Ile and Ser/Thr spectral region. Non-seeded patient fibril spectra and seeded R49G fibril spectra are represented in red and black, respectively. (Right) Superposition of 2D ^15^N, ^13^C correlation spectra focusing on the Cα spectral region. Non-seeded patient fibril spectra and seeded R49G fibril spectra are again represented in red and black, respectively.

## Discussion

Using MAS solid-state NMR, we could identify the residues in FOR005 fibrils that form the rigid core of the fibril. Figure 5 shows a sequence comparison of the FOR005 fibril core with the core identified for other AL fibrils studied to-date either by cryo-EM or MAS solid-state NMR. The experimental amyloidogenic core is indicated in green and yellow for MAS solid-state NMR and cryo-EM experiments, respectively. For the analysis, the proteins AL09 (κ-I, AL patient) (15), MAK33 (κ-IV, murine) (14), AL (λ-I, cardiac AL patient) (12), FOR005 (λ-III, cardiac AL patient), AL55 (λ-VI, cardiac AL patient) (13), and 6aJL2-R24G (λ-VI, model-GL protein) (17) have been employed. Comparison of the different sequences suggests that residues 11-42 and 64-102 are always contained in the amyloidogenic core, involving strands B, CDR1 and C, as well as E, F and CDR3. For fibrils formed by the λ-VI subtype, the N-terminus of the LC is buried in the core (13). This is agreement with a study that suggests that mutations in the N-terminal β-strand accelerate fibril formation (30). Interestingly, the amyloidogenic core for AL55 and 6aJL2-R24G (both λ-VI) are rather similar, suggesting that these two proteins fold into a similar amyloid fibril structure. κ-type sequences, on the other hand, seem to behave differently as the amyloidogenic core involves either the C-terminal or the N-terminal part of the protein sequence. Except for λ-VI, only one EM or NMR study is available for each GL gene segment so far. It remains to be seen whether the fold of the fibril proteins that share a common gene segment are related.

**Figure 5.**
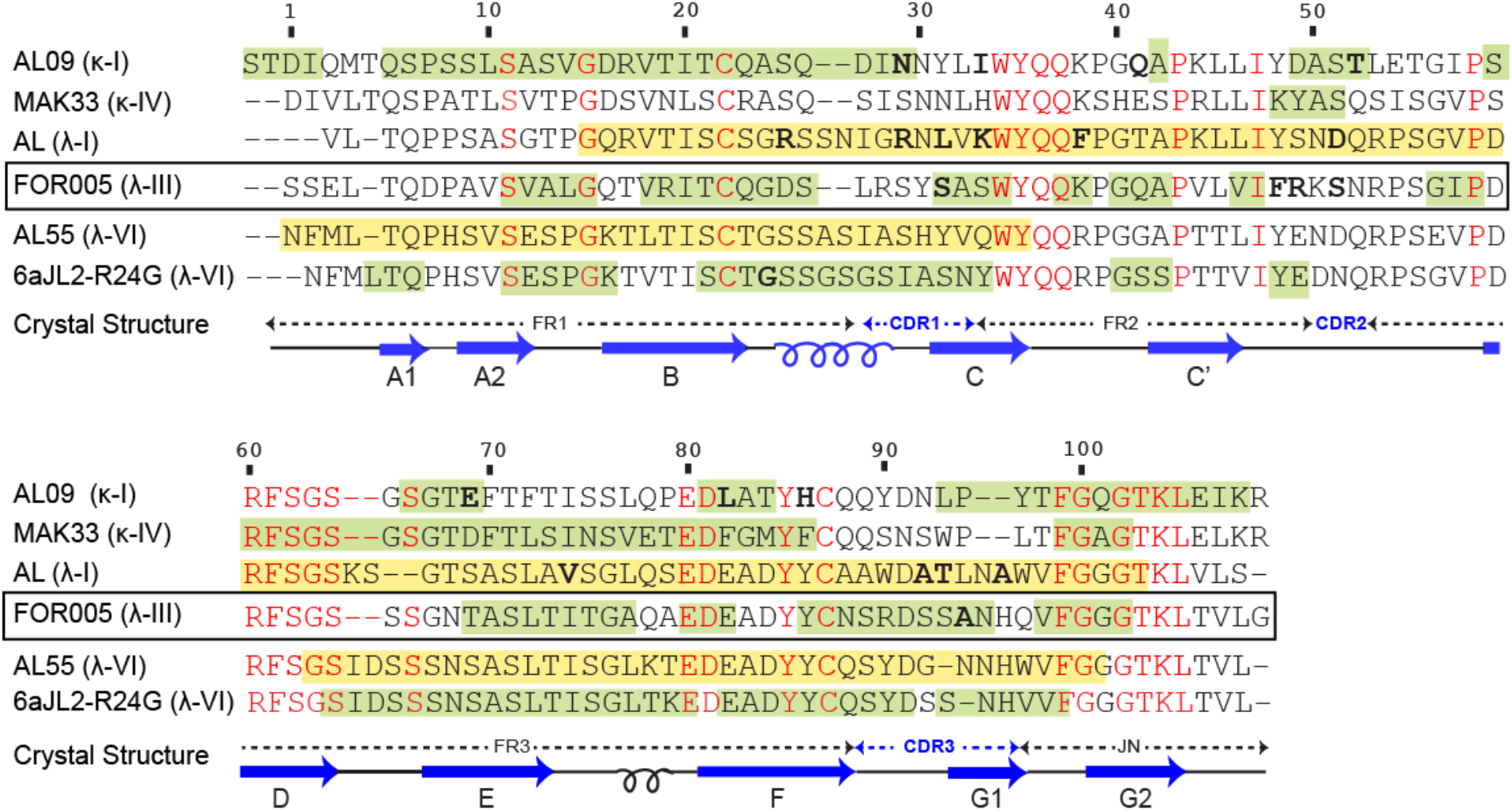
Sequence alignment and amyloidogenic cores of V_L_ fibrils. FOR005 fibrils investigated in this study are highlighted with a black rectangle. For the sequence alignment, amyloid fibril studies involving AL09 (κ-I, AL patient) (15), MAK33 (κ-IV, murine) (14), AL (λ-I, cardiac AL patient) (12), FOR005 (λ-III, cardiac AL patient), AL55 (λ-VI, cardiac AL patient) (13), and 6aJL2-R24G (λ-VI, model-germline protein) (17) are employed. Sequence alignment was performed using the CLUSTALW web tool. The FOR005 primary sequence shares a 40.74 % sequence identity with AL09, 37.38 % with MAK33, 59.81 % for 6aJL2_R25G, 54.72 % with AL, and 57.01 % with AL55, respectively. The amyloidogenic core regions observed by MAS solid-state NMR and cryo-EM are indicated in green and yellow, respectively. Conserved residues among these sequences are highlighted in red. Germline mutations are marked in bold. The secondary structure of the native protein (18) is shown below the sequences in dark blue.

The mechanistic and structural aspects of seeding of amyloid fibrils are not well understood. Linse, Knowles and co-worker have shown that Aβ(1–40) and Aβ(1–42) fibril formation involve a differential amount of primary or secondary nucleation processes (31). Seeding of Aβ(1–40) with Aβ(1–42) does not promote conversion of monomeric Aβ(1–40) into fibrils, although the fibril formation kinetics is accelerated in a concentration dependent manner (32–34). On the other hand, Ramirez-Alvarado and co-worker have reported that LCs can be recruited by homologous and heterologous seeding (27). Furthermore, it has been shown that both AL and MM proteins can be recruited by LC fibrils (35). AL protein recruitment, however, has a much higher efficiency in comparison to recruitment of MM LCs. These findings are in agreement with our observations. We show here that GL protein can be incorporated into V_L_ fibrils using *ex vivo* fibril seeds. The efficiency of fibril formation is, however, reduced. At the same time, we find that the structure of the fibril core is rather similar.

Strikingly, non-seeded FOR005 patient fibrils show very similar spectral patterns as (seeded) R49G or GL fibrils (Figure 4D). In both preparations, many of the ^13^C,^15^N correlation peaks originating from the C-terminus (such as G100, G102) are missing. Seeding in turn stabilizes the C-terminal part of the patient protein in the FOR005 patient fibrils and catalyzes a well-defined adherence of monomeric protein to the fibril core. The single point mutant R49 apparently has a similar effect on fibril structure as seeding, and destabilizes the C-terminus of the protein. This suggests that the template is equally important as the amyloid substrate to form a stable fibril structure.

Interestingly, only two out of the five residues which are mutated in FOR005 with respect to the germline sequence could be assigned to the fibril core. Germline mutations that are part of the amyloidogenic core are S31 and A94 which are located near CDR1 and in CDR3, respectively. On the other hand, the backbone resonances of F48, R49 and S51 in CDR2 could not be assigned, suggesting a high structural heterogeneity for these residues. For R49, only side chain resonances are visible which could be assigned by mutagenesis. The absence of sequential assignments at mutational sites was observed previously for other LCs. For AL09 (κ-I), three out of seven mutated residues could not be assigned, in particular H87Y which was found to be very important for the transition to the altered dimer structure (15,20). Similarly, the λ-I and λ-VI V_L_ proteins investigated by Radamaker et al. (12) and Swuec et al. (13) contain 10 and 12 mutations with respect to the closest germline sequence (S25R, S31R, T33L, N35K, L40F, N53D, I76V, D94A, S95T, G98A; V18L, R24G, G27A, N32H, S43G, S44A, V48L, D52N, N53D, G58E, S96G, S97N). In the cryo-EM structure presented by Swuec et al., 6 out of 12 mutations are occurring in residues 38-65 which are not refined in the structural model (13). In Radamaker et al., all mutations are found in well-defined regions of the electron densities (12). However, no detailed analysis has been performed yet to find out whether a particular residue plays an important role in fibril formation. Two explanations can account for this behavior: First, mutated residues stabilize aggregation intermediate states, and this way catalyze fibril formation. This assumption would be supported by the fact that R49 is solvent exposed in the native state. It seems likely that differences in thermodynamic stability between the native protein and the R49G variant are due to a stabilization of an unfolding intermediate state, which in turn is in agreement with the observation that the fibril formation kinetics is much faster for the patient protein containing the single point mutation R49G in comparison to germline protein. The observed salt bridge cross peaks would thus imply a tertiary contact involving the aggregation intermediate state and not the mature fibril. However, given that the MAS solidstate NMR spectra do not contain any other intermediate state resonances, we do not favor this interpretation. Alternatively, important germline mutations might act as a plug and stabilize the fibril structure in a non-canonical way. As a consequence, only side chain resonances of e.g. R49 are prominently visible in the MAS solid-state NMR spectra, while the backbone resonances are not readily assigned. A detailed analysis of the structural changes between FOR005 patient fibrils and germline fibrils require a structural model with atomic level resolution. Due to sensitivity issues, we were not able to collect a greater number of long-range distance restraints using e.g. PAR or PAIN type experiments so far (36,37). Work into this direction is currently in on-going in our laboratory.

## Conclusion

Our results shed light on the fibril core, polymorphism and the effect of mutations in AL amyloid fibrils. We have identified the core of the amyloid fibrils formed by the patient sequence FOR005. We find that R49 is an important residue that stabilizes the fibril structure via electrostatic interactions. In fibrils formed by protein containing the single point mutation R49G and by the GL sequence, these interactions are lost which yields a destabilization of the C-terminal part of the protein sequence (residues 80-102). By contrast, the N-terminal part of the fibril remains conformationally homogeneous. Analysis of the Cα secondary chemical shifts suggests that R49G fibrils adopt a similar fold as patient protein fibrils. Heterologous seeding experiments indicate that native LC protein can be recruited into pathogenic AL fibrils, which contributes to structural heterogeneity in AL amyloidosis.

## Acknowledgement

This work was performed in the framework of the Research Unit FOR 2969 (German Research Foundation DFG; projects SP01, SP02, SP04, SP04 and SP05). We are grateful to the Center for Integrated Protein Science Munich (CIPS-M) for financial support. We acknowledge support from the Helmholtz-Gemeinschaft.

## Materials

### Source of AL fibrils

AL amyloid fibrils were extracted from the heart of a patient suffering from advanced heart failure due to AL amyloidosis. The AL protein sequence (FOR005) corresponds to “AL case 1” reported by Annamalai et al. (18). Fibrils employed for seeding are extracted from heart tissue as described there and are referred to as *ex-vivo* seeds. The work was conducted on the basis of a valid ethical clearance of the “Ethikkommission der TU München”, project 406/18-AS. Informed consent was obtained from patient FOR005.

### In vitro prepared fibrils

To prepare *in-vitro seeds,* first non-seeded fibrils were prepared. These preformed fibrils were subsequently sonicated for 3 mins, and added to the purified, monomeric protein. This step was repeated two times. In all iterative steps, 5 % w/v seeds were added to monomeric protein to finally select for the fastest growing polymorph.

### Protein expression and purification

Recombinant protein production were purified as described previously (14,38). Briefly, *E. coli* BL21 with a pET28(b+) vector containing the FOR005 gene were grown in minimal medium. Expression was induced with 1 mM IPTG at OD 0.6–0.8. After overnight expression at 37°C, cells were harvested, and inclusion bodies were isolated. The dissolved protein from inclusion bodies was subjected to anion exchange chromatography followed by refolding using a 3.5 kDa dialysis tube and a buffer containing redox agents. Finally, pure protein was obtained using gel filtration chromatography. Total yield was on the order of 15–20 mg protein per liter of culture. To produce isotopically labelled protein, ^15^NH_4_Cl and ^13^C-glucose were employed as nitrogen and carbon sources, respectively. All mutants were purified in the same way as the patient protein.

### CD spectroscopy

CD measurements were performed using a Jasco J-720 spectropolarimeter (Jasco, Grossumstadt, Germany) equipped with a Peltier element. Far-UV CD spectra were measured using 10 μM protein in a 1 mm path length cuvette between 260 and 200 nm. Near-UV CD was measured between 320 and 260 nm using 50 μM protein in a 1-mm cuvette. All spectra were accumulated 16 times, and buffer was corrected. Thermal transitions were recorded using 10 μM protein at 215 nm for VL variants with a heating and cooling rate of 20 °C/h.

### Fibril sample preparation for solid-state NMR

Fibrils were prepared using an initial protein concentration of 50 μM in PBS buffer, pH 6.5 at 37°C. Protein solutions were incubated in a shaker (Thermo Scientific) at 120 rpm. 2.5-5 % seeds were added to yield seeded fibrils. In addition, 0.05 % sodium azide was used to prevent bacterial growth. Samples were incubated for 1 or 2 weeks to yield seeded and non-seeded fibrils, respectively. For all solid-state NMR samples, approx. 15 mg of protein have been employed. Protein aggregates were first centrifuged to reduce the volume to approx. 500 μL. Subsequently, the fibril slurry was sedimented for 1 h into a 3.2 mm thin wall ZrO2 MAS rotor (Bruker, Biospin), using a rotor filling tool (Giotto Biotech) and a L-100 XP ultracentrifuge (Beckman Coulter) equipped with an SW 32 Ti swinging bucket rotor operating at 28.000 rpm. The volume of the MAS rotor has been restricted to the active volume of the NMR coil using teflon spacers.

### Transmission electron Microscopy (TEM)

In order to confirm that fibrils have been formed, we performed TEM experiments. Formvar/Carbon 300 mesh copper coated carbon grids (Electron Microscopy Sciences) was exposed first to an argon atmosphere for 10 s. 5 μL of sample was then added to the grids and incubated for 1 minute. Grids were subsequently washed with water and dried in filter paper. For staining, 10 μl of uranyl acetate (2 %) was added for up to 30 s. Extra stain was removed from the grid using filter paper. Grids were visualized in TEM employing a Zeiss EM 10 CR or a LIBRA 120 plus microscope (Zeiss, Germany).

### ThT kinetics assay

Fibril formation kinetics was monitored by standard ThT assay (27,39,40). Triplicates were performed for all samples. 0.02 % sodium azide was added to avoid bacterial growth. The 50 μM V_L_ protein samples (with or without 2.5 % - 5 % seeds) were incubated with 25 μM ThT, using a 96 well plate (Thermofisher scientific) and measured in fluorescence spectrometer (PHERAstar plus, BMG LABTECH) with fluorescence excitation and emission wavelength of 440 nm and 480 nm, respectively. The ThT experiments have been carried out at a temperature of 37 °C. During incubation, the samples were agitated at 500 rpm using thermoshaker (PST-60HL-4, Biosan).

### Solid state NMR experiments

All solid-state NMR experiments are carried out at an external magnetic field of 17.6 T (corresponding to a ^1^H Larmor frequency of 750 MHz). 2D ^13^C,^13^C correlation experiments were acquired using either PDSD or DARR for mixing. Experiments involving aliphatic carbons were performed at a MAS frequency of 10 kHz with using a ^13^C,^13^C missing time of 50 ms. Experiments involving aromatic residues were performed at a MAS frequency of 16.5 kHz to avoid interference with rotation side bands. To assign the fibril NMR chemical shifts, conventional 3D NCACX and 3D NCOCX were recorded (41,42). For ^13^C,^15^N transfers, specific CP based experiments were employed (43). In addition, 3D CONCA and 3D CANCO experiments were performed to confirm and assign ambiguous residues (44,45). In these experiments, optimal control CP (OC-CP) were used to gain sensitivity (46). The effective sample temperature was adjusted to 0 °C. In order to characterize salt bridges, 2D ^13^C,^15^N TEDOR experiments (28,47) have been recorded. In these experiments, the MAS rotation frequency has been adjusted to 16.5 kHz MAS, using short (1.9 ms) and long (15.0 ms) TEDOR mixing times. In 3D experiments, 25 % NUS (48) was used to gain sensitivity and to reduce experimental time. NUS spectra were reconstructed using the mdd algorithm in TOPSPIN employing the NUS plugin (49). The secondary chemical shifts were calculated according to the formula: [Cα (observed) – Cα(random coil)] – [Cβ(observed) – Cβ(random coil)].

